# Network Analysis of the Cerebrospinal Fluid Proteome Reveals Shared and Unique Differences Between Sporadic and Familial Forms of Amyotrophic Lateral Sclerosis

**DOI:** 10.1101/2024.02.29.582840

**Authors:** Adam N. Trautwig, Edward J. Fox, Eric B. Dammer, Anantharaman Shantaraman, Lingyan Ping, Duc M. Duong, Allan I. Levey, James J. Lah, Christina N. Fournier, Zachary T. McEachin, Jonathan D. Glass, Nicholas T. Seyfried

## Abstract

**Background:** Amyotrophic Lateral Sclerosis (ALS), a neurodegenerative disease involving loss of motor neurons, typically results in death within 3-5 years of disease onset. Although roughly 10 % of cases can be linked to a specific inherited mutation (e.g., C9orf72 hexanucleotide repeat expansion or SOD1 mutation), the cause of the majority of cases is unknown. Consequently, there is a critical need for biomarkers that reflect disease onset and progression across ALS subgroups.

**Methods:** We employed tandem mass tag mass spectrometry (TMT-MS) based proteomics on cerebrospinal fluid (CSF) to identify and quantify 2105 proteins from ALS patients with sporadic disease (n=35), C9orf72 ALS (n=10), and SOD1 ALS (n=6), as well as age-matched healthy controls (n=44) and asymptomatic C9orf72 carriers (n=6). We used differential protein abundance and network analyses to determine how protein profiles vary across disease types in ALS CSF.

**Results:** Integrated differential and co-expression network analysis identified proteomic differences between ALS and control, and differentially abundant proteins between sporadic, C9orf72 and SOD1 ALS. Groups of proteins also differentiated asymptomatic C9orf72 mutation carriers from those with C9orf72 ALS, marking a pre-symptomatic proteomic signature of C9orf72 ALS. Similarly, additional proteins differentiated asymptomatic from controls. Leveraging additional publicly available ALS and AD proteomic datasets, we validated our ALS CSF network and identified ALS-specific proteins within Module 5 (M5)-Extracellular matrix (e.g., IGF2, RARRES2, LGALS3, GALNT15, and LYZ) and shared biomarkers across neurodegenerative diseases linked to Module 10 (M10)-Ubiquitination/Gluconeogenesis (e.g., NEFL, NEFM, CHIT1, and CHI3L1).

**Conclusions:** This study represents a comprehensive analysis of the CSF proteome across sporadic and genetic causes of ALS that resolves differences among these disease subgroups and points to varying pathogenic pathways that result in disease.

## Introduction

Amyotrophic Lateral Sclerosis (ALS) is a heterogeneous motor neuron disease that typically results in death within 3-5 years (1, 2). Clinical manifestations include a spectrum of upper and lower motor neuron involvement and wide variability in disease progression. Roughly 10% of ALS cases are driven by an inherited mutation, of which the most common are C9orf72 hexanucleotide repeat expansion (3, 4) and point mutations in the gene for superoxide dismutase 1 (SOD1) (5). Though the pathogenic mechanisms underlying the various genetic forms of ALS are likely to be distinct from sporadic disease, the clinical presentations are remarkably similar, making the discovery of biomarkers that distinguish the various forms of ALS of paramount importance. Also, in people who harbor disease-causing mutations but remain asymptomatic, comparison of biomarkers from the pre- and post-symptomatic phase will provide insight into potential markers of disease transition, and also mark an early time point when disease modifying therapies could be started. A focus on cerebrospinal fluid (CSF) biomarkers allows for interrogation of CNS protein changes that may differentiate disease pathways among various pathogenic forms of ALS as well as provide tools allowing for early diagnosis and monitoring of disease activity.

Here, our objective was to evaluate the CSF proteome in order to enhance our comprehension of both shared and distinct disease alterations associated with CNS cell-types and pathways across sporadic and genetic ALS subgroups.

Mass spectrometry-based proteomics coupled with systems biology approaches using co-expression network analysis is a valuable tool for discovery of disease biomarkers and pathways, including in ALS and Alzheimer’s Disease (6–10). Unbiased proteomics of human brain and CSF coupled with network analysis has emerged as a valuable tool for organizing proteome-wide expression data into groups or “modules” of highly correlated proteins that reflect various biological functions linked to neurodegeneration (11). While ALS brain proteomic networks have been examined (6), ALS CSF proteomic networks from large cohorts that include both sporadic and familial ALS across different mutation carriers have been under-investigated.

To this end, we performed unbiased CSF proteomics from sporadic ALS (sALS), C9orf72 ALS, C9orf72 asymptomatic carriers, SOD1 ALS, and healthy controls. This cohort enabled measurement of differentially abundant proteins (DAPs) between i) each ALS subgroup and control; ii) symptomatic and asymptomatic C9orf72; and iii) genetic and sporadic ALS. We also validated changes in the CSF proteome from the Emory ALS cohort with a published, independent ALS CSF proteomic dataset (12). In addition, to address disease specificity, we compared potential biomarkers in these ALS cohorts to Alzheimer’s disease (AD) CSF. Network modules found to change significantly with ALS type were associated with Module 5 (M5)- Extracellular matrix/Heparin binding, M7-Cytoskeleton/Microglia, and M10- Ubiquitination/Gluconeogenesis. DAPs, relative to controls, were identified in each group. Of particular interest were proteins that differentiated C9orf72 ALS from asymptomatic C9orf72 carriers that may be used to identify a transition from asymptomatic to symptomatic disease. Collectively our findings suggest that while sporadic and genetic forms of ALS display largely overlapping CSF proteomes, differences point to unique pathogenic pathways.

## Materials and Methods

### CSF samples

All CSF samples were collected as part of ongoing research at the Emory ALS Center in Atlanta, Georgia. Research participants provided informed consent under protocols approved by the Institutional Review Board at Emory University. The Emory ALS CSF cohort contained samples from C9orf72 ALS (n=10), C9orf72 asymptomatic carriers (n=6), SOD1 ALS (n=6), and sALS (n=35), as well as age matched healthy controls (n=44). Characteristics of the Oh et al. ALS (12) and Emory AD (disease specificity) CSF cohorts were previously published (13). Characteristics of our Emory ALS cohort are summarized in **Table 1** and detailed in **Supplemental Table 1**.

**Table 1.** Characteristics of the Emory ALS cohort (i.e. Set 1). Unabridged cohort traits are enumerated in Supplemental Table 1. Abbreviations F=Female, M=Male; AA=African American, A=Asian, C=Caucasian, H/L=Hispanic/Latino; B=Bulbar, LE=Lower Extremity, U=Unknown, UE=Upper Extremity, y=Years, mo=Months. “*” 10/10 C9orf72 ALS deceased, 19/34 sALS deceased. 2/6 SOD1 ALS deceased, average not calculated.

|  | **Control** | **C9orf72 Asymptomatic** | **C9orf72 Positive** | **SOD1 Positive** | **Sporadic ALS** |
| --- | --- | --- | --- | --- | --- |
| **Sample number (n)** | 44 | 6 | 10 | 6 | 35 |
| **Diagnosis** | 44 HC | 6 C9-asym | 10 C9 | 6 SOD1 | 29 sALS, 6 PLS |
| **Site onset** | - | - | 5 B, 3 LE, 2 UE | 3 LE, 2 U, 1 UE | 3 B, 17 LE, 15 UE |
| **Sex** | 27 F, 17 M | 2 F, 4 M | 4 F, 6 M | 2 F, 4 M | 13 F, 22 M |
| **Race** | 44 C | 5 C, 1H/L | 10 C | 6 C | 3 AA, 1 A, 31 C |
| **Age at Onset (y)** | - | - | 56.4 (±8.9) | 58.3 (±4.7) | 53.9 (±11.5) |
| **Age at Sample (y)** | 64.1 (±7.6) | 51.5 (±18.5) | 57.2 (±8.7) | 59.3 (±4.6) | 56.7 (±10.9) |
| **Duration of disease (mo)\*** | - | - | 41 (+/- 26) | - | 38 (+/- 26) |

### Protein digestion and Tandem Mass Tag (TMT) labeling of CSF

In order to sample the CSF in an unbiased manner and given that we have previously shown that immunodepletion resulted in only a marginal improvement in proteomic coverage, the CSF samples were not immunodepleted prior to digestion (14–16). First, 50 μL of CSF was transferred to 1 mL deep well plates for digestion with lysyl endopeptidase (LysC) and trypsin. The samples were then reduced and alkylated with 1 μL of 0.5 M tris-2(-carboxyethyl)- phosphine (ThermoFisher) and 5 μL of 0.4 M chloroacetamide in a 90 °C water bath for 10 min followed with 5 min bath sonication. After the sample was cooled down on ice, 56 μL of 8 M urea buffer (8 M urea, 10 mM Tris, 100 mM NaH_2_PO_4_, pH 8.5) with 12.5 mAU of LysC (Wako), was added to each sample, resulting in a final urea concentration of 4 M.

Samples were then mixed well, gently spun down, and incubated overnight at 25 °C for digestion with LysC. The following day, samples were diluted to 1 M urea with a mixture of 360 μL of 50 mM ammonium bicarbonate (17) and 5 μg of Trypsin (ThermoFisher). The samples were subsequently incubated overnight at 25 °C for digestion with trypsin. The next day, the digested peptides were acidified to a final concentration of 1% formic acid and 0.1% trifluoroacetic acid. This was immediately followed by desalting on 30 mg HLB columns (Waters) and then eluted with 1 mL of 50% acetonitrile (ACN) as previously described (18). To normalize protein quantification across batches 150 μl of elution was taken from all CSF samples and then combined to generate a pooled sample as previously described (18). This pooled sample was split into 850 μL each as global internal standards (GIS) (19). All individual samples and the GIS standards were then dried using a speed vacuum. Six TMT batches were balanced for diagnosis, age, and sex using ARTS (automated randomization of multiple traits for study design) (20). Using an 18-plex Tandem Mass Tag (TMT-pro) kit (ThermoFisher, Lot# UK297033 and WI336758), 17 channels were allocated for CSF samples with the remaining channel (126) containing a GIS pool, as described in (16).

### High-pH peptide fractionation

Dried samples were re-suspended in high pH loading buffer (0.07% vol/vol NH4OH, 0.045% vol/vol FA, 2% vol/vol ACN) and loaded onto a Waters BEH column (2.1 mm × 150 mm with 1.7 µm particles). A Vanquish UPLC system (ThermoFisher Scientific) was used to carry out the fractionation. Solvent A consisted of 0.0175% (vol/vol) NH4OH, 0.01125% (vol/vol) FA, and 2% (vol/vol) ACN; solvent B consisted of 0.0175% (vol/vol) NH_4_OH, 0.01125% (vol/vol) FA, and 90% (vol/vol) ACN. The sample elution was performed over a 25 min gradient with a flow rate of 0.6 mL/min with a gradient from 0 to 50% solvent B. A total of 96 individual equal volume fractions were collected across the gradient. Fractions were concatenated to 48 fractions and dried to completeness using vacuum centrifugation.

### Mass-spectrometry analysis and data acquisition

All samples (∼ 1 µg for each fraction) were loaded and eluted by an Ultimate 3000 RSLCnano (Thermo Scientific) with an in-house packed 20 cm, 150 μm i.d. capillary column with 1.7 μm CSH (Water’s) over a 22 min gradient. Mass spectrometry was performed with a high-field asymmetric waveform ion mobility spectrometry (FAIMS) Pro front-end equipped Orbitrap Eclipse (Thermo) in positive ion mode using data-dependent acquisition with 1.5 s top speed cycles for each FAIMS compensative voltage. Each cycle consisted of one full MS scan followed by as many MS/MS events that could fit within the given 1 s cycle time limit. MS scans were collected at a resolution of 120,000 (410–1600 m/z range, 4 × 10^5 AGC, 50 ms maximum ion injection time, FAIMS compensative voltage of -45 and -65). Only precursors with charge states between 2 + and 6+ were selected for MS/MS. All higher energy collision-induced dissociation (HCD) MS/MS spectra were acquired at a resolution of 30,000 (0.7 m/z isolation width, 35% collision energy, 1 × 10^5 AGC target, 54 ms maximum ion time, turboTMT on). Dynamic exclusion was set to exclude previously sequenced peaks for 20 s within a 10-ppm isolation window.

### Database search and protein quantification

Database searches and protein quantification was performed on 576 RAW files (96 RAW files/fractions per batch) using FragPipe (version 19.1). The FragPipe pipeline relies on MSFragger (version 3.7) (21, 22) for peptide identification, MSBooster (23), and Percolator (24) for FDR filtering and downstream processing. MS/MS spectra were searched against all canonical Human proteins downloaded from Uniprot (20,402; accessed 02/11/2019), as well as common contaminants (51 total), and all reverse sequences (20,453). The workflow we used in FragPipe followed default TMT-18 plex (i.e., TMTpro) parameters. Briefly, precursor mass tolerance was -20 to 20 ppm, fragment mass tolerance of 20 ppm, mass calibration and parameter optimization were selected, and isotope error was set to -1/0/1/2/3. Enzyme specificity was set to strict-trypsin with up to two missing trypsin cleavages allowed. Cleavage was set to semi Peptide length was allowed to range from 7 to 35 and peptide mass from either 200 to 5,000 Da. Variable modifications that were allowed in our search included: oxidation on methionine, N-terminal acetylation on protein and peptide, TMT labeling reagent modifications on serine, threonine, and histidine with a maximum of 3 variable modifications per peptide (25). The false discovery rate (FDR) threshold was set to 1%. A total of 49,762 peptides which mapped to 2,568 proteins were detected. After filtering out proteins that were absent in 50% or more of specimens 23,743 peptides and 2,105 proteins were retained. We had one dataset of Emory ALS peptides and one dataset of Emory ALS proteins as well as two other previously published datasets. The ALS dataset from Oh et al. (12) (1831 proteins; **Supplemental Table 3**) and the Emory Alzheimer’s disease (disease specificity) dataset (13) (2146 proteins; **Supplemental Table 4**) were retrieved and processed in FragPipe as above, for proteins only, with default parameters for 11 and 16 plex TMT, respectively. We did not process Oh et al. ALS or Emory AD peptides.

### Bioinformatics processing and statistical analysis

We employed a Tunable Approach for Median Polish of Ratio (TAMPOR) (26) and removal of peptides or proteins absent in 50% of cases or greater, as previously published (16, 27, 28). To ensure the reliability of our data, we initiated the analysis by identifying and removing potential outliers, although none were present. We then performed parallelized (29) ordinary, nonparametric, bootstrapping regression to remove variation due to age, sex, and batch effect using an established pipeline (7, 28). Fast parallel one-way ANOVA with Benjamini-Hochberg correction for multiple comparisons was conducted within each disease group using an in-house script (https://github.com/edammer/parANOVA) to identify peptides and proteins that were differentially expressed. Differential abundance is presented as volcano plots, which were generated with the ggplot2 package (30).

### Protein Network Analysis

Weighted Gene Co-expression network analysis (WGCNA) (31) was used to construct modules of co-expressing proteins as previously published (7, 14). Briefly, the blockwiseModules function from the WGCNA package in R was utilized with the following parameters for Emory ALS CSF samples: soft threshold power beta = 4, deepSplit = 4, minimum module size = 15, merge cut height = 0.07, and a signed network with partitioning around medoids (mapping a distance matrix to k clusters, where K is data-adaptively selected) (32). Module correlation to disease type was evaluated with biweight midcorrelation (BiCor) analysis by separating each disease control combination. Fisher’s exact test (FET) was performed for each module’s members against the merged human brain cell type marker list to determine cell type enrichment using an in-house script (https://github.com/edammer/cellTypeFET). Similarly, to determine module gene ontology (GO) a FET was performed for each module member against the Bader Lab’s GMT formatted ontology lists from February 8, 2023 (33) (https://github.com/edammer/GOparallel). One way ANOVA was conducted across all disease type groups for each module eigenprotein.

Module preservation of the Emory ALS dataset in the quantitative data of Oh et al. (12) was assessed using the modulePreservation function of WGCNA with 500 permutations (7, 28, 34). Synthetic module eigenproteins (MEs) were calculated using the top 20 percent of hubs ranked by kME_intramodule_ in the Emory data set used to build the template network, and a minimum of 4 such hubs found in the mapped data set (non-Emory CSF data). The first principal component of the selected proteins in the mapped data set was calculated as the synthetic ME in that data using the moduleEigengenes function of WGCNA R package as in (7, 28, 34).

## Results

### Experimental workflow identifies differentially expressed proteins across disease subgroups

We compared CSF proteomes from sALS (n=35), C9orf72 ALS (n=10), C9orf72 asymptomatic carriers (n=6), SOD1 ALS (n=6), and healthy controls (n=44) with the goals of discovering differences between ALS and control, and also protein signatures that may differentiate the ALS groups (**Figure 1a**). Following QC, TMT-MS proteomic analysis represented 2,105 proteins with more than 50 percent of all samples having quantification. Protein abundance was adjusted for batch, age, and sex (14, 16). As expected, protein levels of neurofilaments (NEFM and NEFL) were increased in both sporadic and genetic ALS samples compared to controls, consistent with neurodegeneration (35–37). Furthermore, we observed an increase in chitinases (CHIT1) and CHI3L1 linked to inflammation and previously shown to increase in ALS (38, 39).

**Figure 1.**
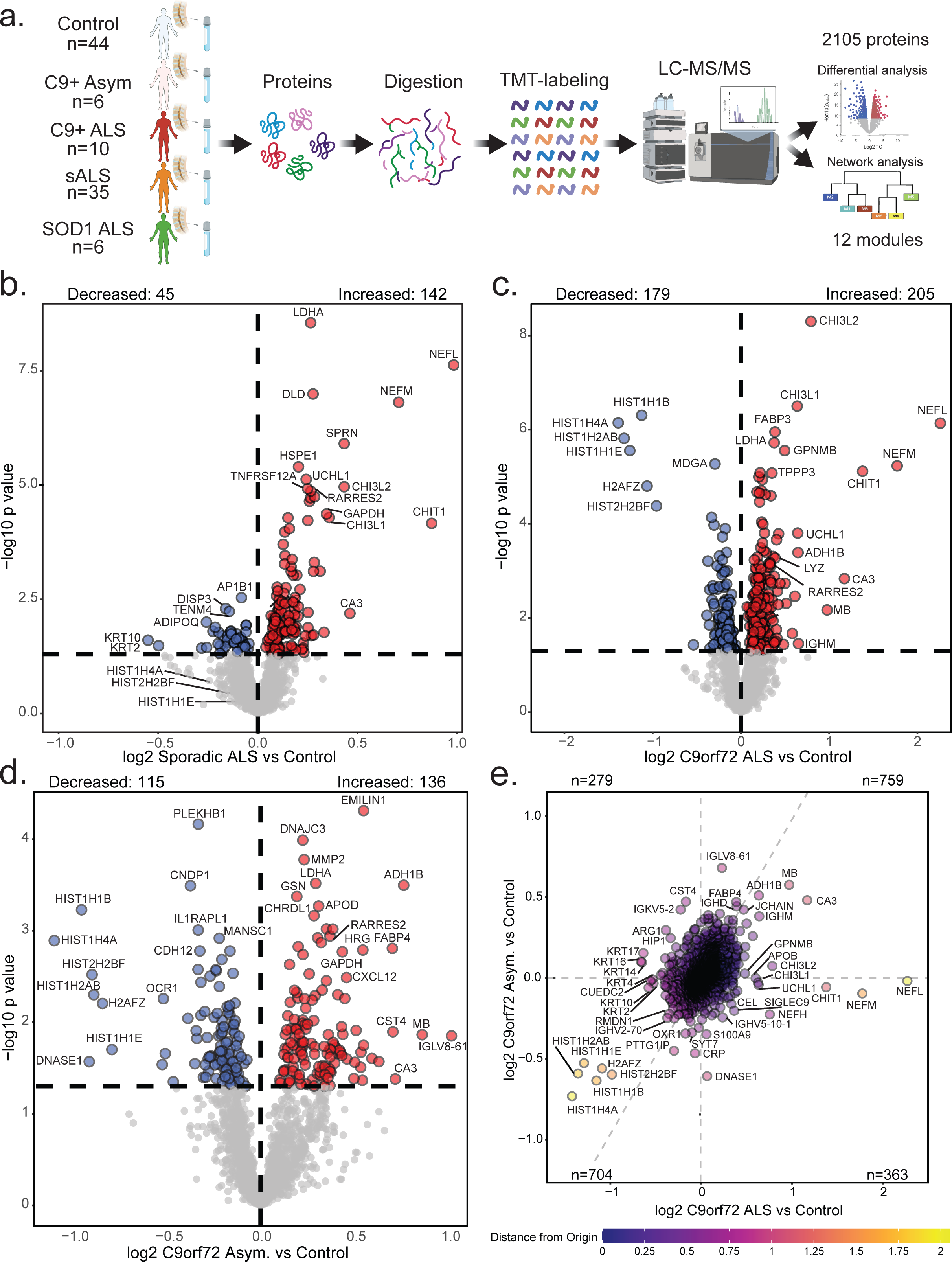
Experimental workflow with differential expression of ALS versus control CSF proteomes. **a.** Schematic of experimental workflow to examine proteomic differences between subjects with ALS and controls in cerebrospinal fluid (CSF). **b-d.** Volcano plots showing differential abundance profiles comparing control CSF (n=44) to that from sALS (n=35), C9orf72 ALS (n=10) and asymptomatic C9orf72 carriers (n=6). Log_2_ fold change (x-axis) and one-way ANOVA with Benjamini-Hochberg corrected by disease -log10 p-values (y-axis). Note the commonly increased CSF proteins in patients with ALS such as NEFL, NEFM, CHIT1, and GPNMB. Proteins significantly (p<0.05) increased in abundance are depicted in red, significantly decreased in blue, and neither in grey. **e**. Scatter plot showing differential expression of C9orf72 ALS versus asymptomatic C9orf72 carriers. Log_2_ fold change of C9orf72- Associated ALS (x-axis) and Log_2_ fold change of C9orf72 Asymptomatic carriers (y-axis) were compared for each protein (n=2105). Proteins are colored based on distance from the origin.

Comparison of CSF samples between sporadic and C9orf72 ALS demonstrated increases in many of the same proteins as compared to controls (**Figures 1b and 1c**). These increased proteins included proteins with axonal regeneration ontology as well as proteins with metabolic pathways associated with L-ascorbic acid, cellular ketone, and cellular aldehyde. Proteins related to chemical synaptic transmission and assembly of postsynaptic elements in C9orf72 ALS CSF were significantly decreased in abundance relative to sALS.

Analysis of CSF from asymptomatic C9orf72 carriers found 251 differentially abundant proteins (DAPs) compared to controls (**Figure 1d**). In comparison to those with C9orf72 ALS, asymptomatic mutation carriers showed reduced abundance of proteins related to amino sugar catabolism and keratin binding (**Figure 1e**). Conversely, C9orf72 mutation carriers with ALS showed reduced abundance of proteins associated with keratinization, keratinocyte differentiation, and intermediate filament organization as compared to asymptomatic C9orf72 carriers. Many of the protein differences become more significant in symptomatic C9orf72 ALS, suggesting that the C9orf72 repeat expansion induces protein change in pre-symptomatic stages of disease, and these changes continue during progression into the symptomatic phase.

SOD1 protein was surprisingly decreased in abundance in CSF from SOD1 patients compared to all other forms of ALS as well as compared to controls (**Figure 2a; Supplemental Table 2,5**). Total SOD1 abundance assembled from peptide-level data was also analyzed to assess whether the observed decrease in SOD1 abundance was representative across the protein sequence and not solely driven by peptides which overlap with the single-amino acid variants caused by the familial mutations (**Figure 2a)**. Consistent with the volcano plot, total SOD1 (all summed peptides measured) was decreased in abundance across all SOD1 mutation carriers compared to controls and other ALS subgroups, with A5T patients demonstrating the lowest levels of SOD1 (**Figure 2b)**. Fully-tryptic peptides from all 6 SOD1 carriers covered 64.9% of the SOD1 protein (**Figure 2c**) and in all but one peptide, a significant decreased in SOD1 peptide abundance was determined in SOD1 patients regardless of whether the peptide overlapped with a variant caused by the mutation. Consistent with the total SOD1 levels, SOD1 carriers with mutation A5T had the lowest peptide abundance levels across the primary sequence (**Figure 2d**). In addition to the reduction in SOD1, several histones were also decreased in abundance in SOD1 ALS patients, a difference also identified in C9orf72 mutation carriers (**Supplemental Table 2**) suggesting a shared mechanism in histone dysregulation with C9orf72 and SOD1 mutant carriers. In summary, the protein biomarkers we identified are associated with each type of ALS, genetic versus sporadic causes of ALS, as well as proteins that may indicate a shift from asymptomatic to symptomatic disease states.

**Figure 2.**
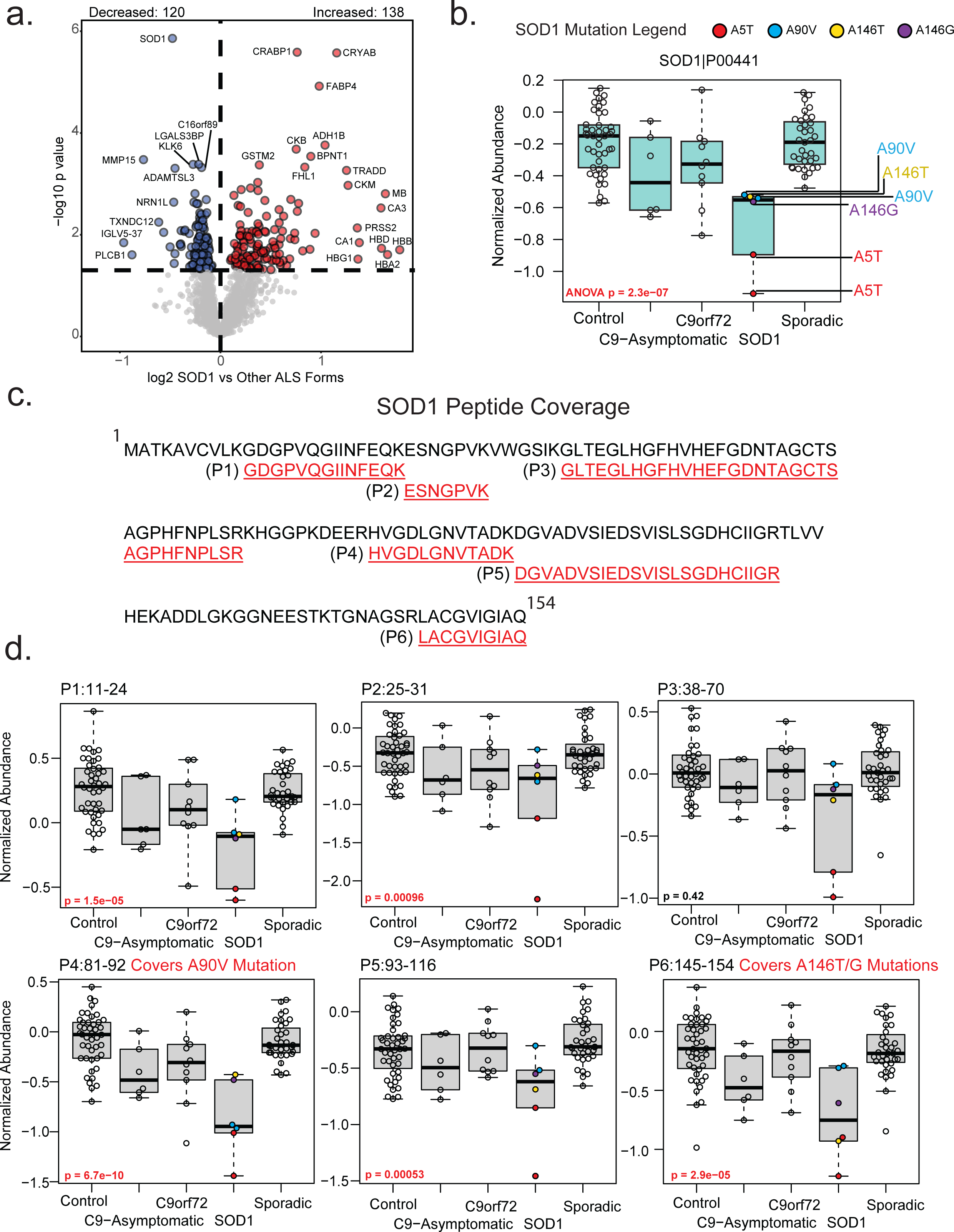
SOD1 protein and peptide-specific CSF levels for SOD1 mutant ALS carriers. **a.** Volcano plot representing differential protein abundance comparing SOD1-associated ALS cases (n=6) versus all other ALS cases (n=45). Log_2_ fold change (x-axis) and one-way ANOVA with Benjamini-Hochberg corrected by disease -log10 p-values (y-axis) are shown for each protein (n=2,105). Proteins that were significantly (p≤0.05) down in disease (SOD1 ALS, relative to all other ALS) are depicted in blue (n=120). Proteins that were significantly up are depicted in red (n=138), and proteins that were neither significantly up nor down are grey. **b.** Total SOD1 normalized protein abundance levels across all subgroups. One-way ANOVA was used to determine if a difference was present between the ALS groups. Individual SOD1 mutations are depicted by color. **c.** SOD1 specific-peptide level quantification across controls and disease subgroups. Peptides were visualized for overlap of the canonical SOD1 protein sequence (P00441). Only fully-tryptic peptides that were detected in all 6 SOD1 ALS patients were included. **d.** Boxplots for abundance of each peptide identified were evaluated with one-way ANOVA. Each datapoint in the SOD1 ALS group is annotated with the mutation associated with each patient. Peptides that overlap with SOD1 regions that have mutations (single amino acid substitutions) in our cohort are emphasized in red with the specific mutation noted.

### Network analysis of the ALS CSF proteome reveals modules related to brain pathways, brain cell-types and genetic background

We utilized Weighted Gene Co-expression Network Analysis (WGCNA) (31) to identify modules or ‘communities’ of proteins that are highly correlated with across CSF samples. These modules in CSF reflect various biological functions linked to brain cell types and ontologies (15, 16). Using the first principal component of all proteins in a module, or the module eigenprotein, we can relate the abundance of each module to disease phenotypes with greatly reduced reliance on multiple testing.

Here, our network modules ranged from 476 (M1) to 29 (M12) member proteins with 409 proteins not nesting into a module. The 12 network modules were generally divided into three branches of relatedness, allowing us to infer which modules were most similar (**Figure 3a**). We compared correlation between specific disease cohorts and network module co-expression to determine that five modules were associated with C9orf72, three with SOD1, and two with sALS, with overlap in module identity. We also compared module membership (**Supplemental Table 6**) and proteins associated with specific brain cell-types for overlap in order to ascertain overarching trends driven by shifts across individuals in brain processes reflecting relative cell type abundance or activity due to disease or interindividual variation. Three modules had significant overlap with neurons (M1, M4, and M12) and microglia (M7, M8, and M9), two with oligodendrocytes (M1 and M4), and one with endothelia (M2; **Figure 3b**). Modules showing a high degree of correlation did not necessarily have more overlap with protein markers of cell type. Specifically, M1 and M4 were significantly enriched with neuronal and oligodendrocyte associated proteins but were more closely correlated to modules nine and 12; respectively (**Figure 3c**). The three largest modules corresponded to M1- Neuronal, M2-Complement activation enriched with markers specific to endothelia, and M3-Adaptive immune response ontologies (**Figure 3d**). Smaller modules were associated with M4-Neuron development, M5- Extracellular matrix/Heparin binding, M6-Lysosomal/Vesicle, M7-Cytoskeleton/Microglial, M8- Inflammatory Response, M9-Lysosome, M10-Ubiquitination/Gluconeogenesis, M11- Postsynaptic membrane/Signaling, and M12-Nervous system development (**Supplemental Table 7)**. Modules associated with M5-Extracellular matrix/Heparin binding (p=0.0084), M7- Cytoskeleton/Microglia (p=0.025), and M10-Ubiquitination/Gluconeogenesis (p=2.4e-07) varied significantly among control and ALS groups (**Figure 4a**). To reinforce these findings, most of the increased DAPs in ALS cases irrespective of genetic cause mapped to M7, M5 and M10, whereas decreased DAPs in ALS were distributed in M4, M11, and M12 (**Figure 4b**). DAPs in C9orf72 patients increased in abundance also contributed to these modules, with lowered abundance proteins belonging to modules associated with M4-Neuron development and M12-Nervous system development. Proteins increased in abundance in asymptomatic C9orf72 cases were more commonly clustered into the module associated with M5-Extracellular matrix/Heparin binding and proteins that were significantly increased in C9orf72 ALS relative to sporadic cases, including neurofilaments, chitinases, and deubiquitinases belonging to the module associated with M10-Ubiquitination/Gluconeogenesis (**Figure 4b**). Module overarching ontologies and correlation helped identify broad protein functions more closely associated with C9orf72 (M5-Extracellular matrix/Heparin binding) and SOD1 (M8-Inflammatory Response) as well as the degree to which these functions overlapped (i.e. all ALS subgroups correlated to M7- Cytoskeleton/Microglial and M10-Ubiquitination/Gluconeogenesis). Overall, network analysis effectively organizes the CSF proteome into protein modules that are strongly linked to hallmark ALS and neurodegenerative biomarkers, including NEFL.

**Figure 3.**
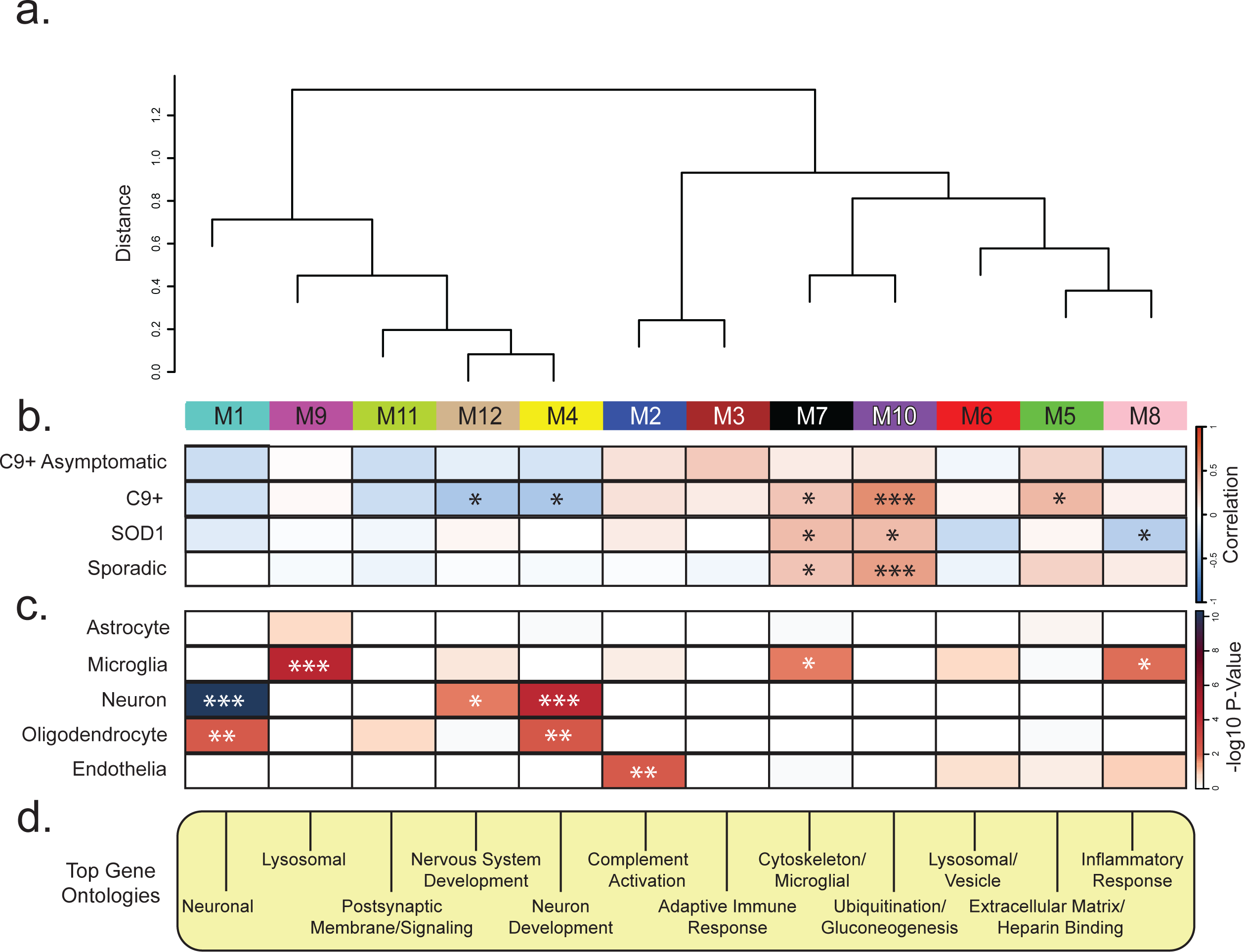
CSF Network modules associate with brain cell-types ALS disease subtypes. **a.** Cluster dendrogram indicates similarity of WGCNA network modules based on correlation of eigenproteins (i.e., first principal component). **b.** Relationship between ALS disease subgroup (sporadic, SOD1, Asymptomatic C9orf72 and symptomatic C9orf72) with individual protein modules was evaluated by cross referenced trait values with module proteins using a Biweight midcorrelation (BiCor) analysis. Significance as determined by BiCor are denoted by overlain asterisks; \**p* < 0.05, \*\**p* < 0.01, \*\*\**p* < 0.001. Note the relatedness of modules 7 and 10 as well as the overlap in significance between these modules by disease subtype compared to control group. **c.** Cell-type enrichment was characterized by comparing module proteins with a list of proteins known to be enriched in astrocytes, microglia, neurons, oligodendrocytes, and endothelia; respectively (see methods). Significance levels determined by one-tailed Fisher’s exact test are denoted by overlain asterisks; \**p* < 0.05, \*\**p* < 0.01, \*\*\**p* < 0.001. **d.** Top gene ontology (GO) terms were selected from significant GO annotations (**Supplemental Table 7)**.

**Figure 4.**
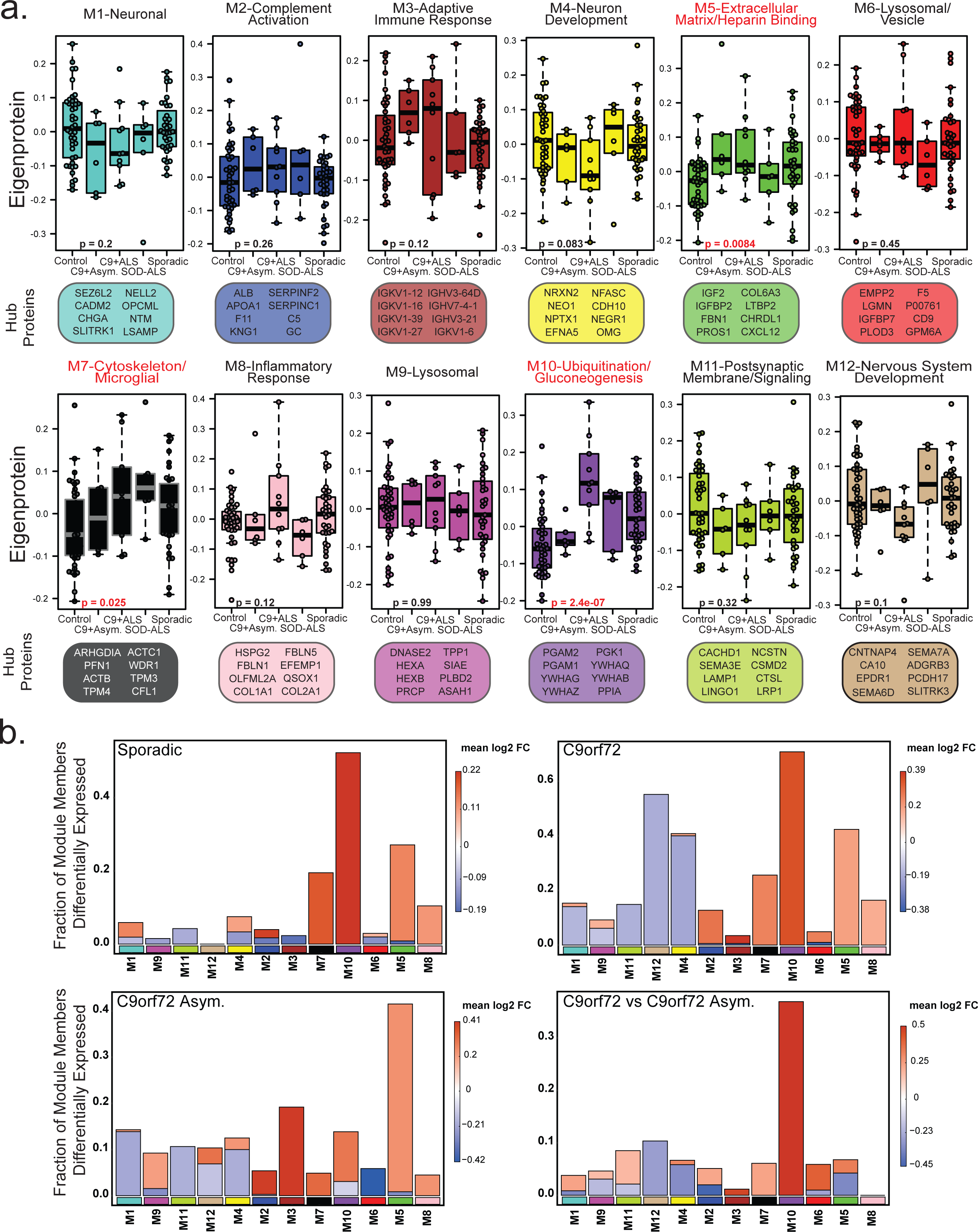
CSF Network Modules vary across ALS subgroups. **a.** Eigenproteins for each CSF proteome modules (n=12) were compared across control and ALS disease type by one-way ANOVA. Hub proteins and GO terms for each module are highlighted **b.** Differentially abundant proteins from each subgroup compared to controls for C9 ALS, Sporadic ALS and C9 Asymptomatic were mapped by module. Symptomatic vs asymptomatic C9orf72 differences were also included. The height of the bars represents the fraction of module member proteins that were differentially abundant. The bars are color coded by heatmap for average log_2_ difference in abundance, where red represents an increase in abundance, and blue represents a decrease in abundance.

## Validation of protein abundance changes in an independent cohort of CSF

To assess the consistency of our findings in the Emory ALS CSF, we analyzed a TMT-MS dataset from a second independent ALS CSF proteomic dataset ALS CSF proteome (12), allowing us to compare the findings of our analysis of individual proteins as well as co-expressing modules in an independent cohort. Hereafter, the Oh et al. ALS dataset will be referred to as “Set 2”. The aim was to view these data collectively, where Set 1 (Emory ALS) was generated using TMTpro reagents (TMT18) and Set 2 generated using the first generation TMT11 reagents. Set 1 ALS CSF proteomics was compared to Set 2 at both the individual protein and network module level, as outlined in **Figure 5a**.

**Figure 5.**
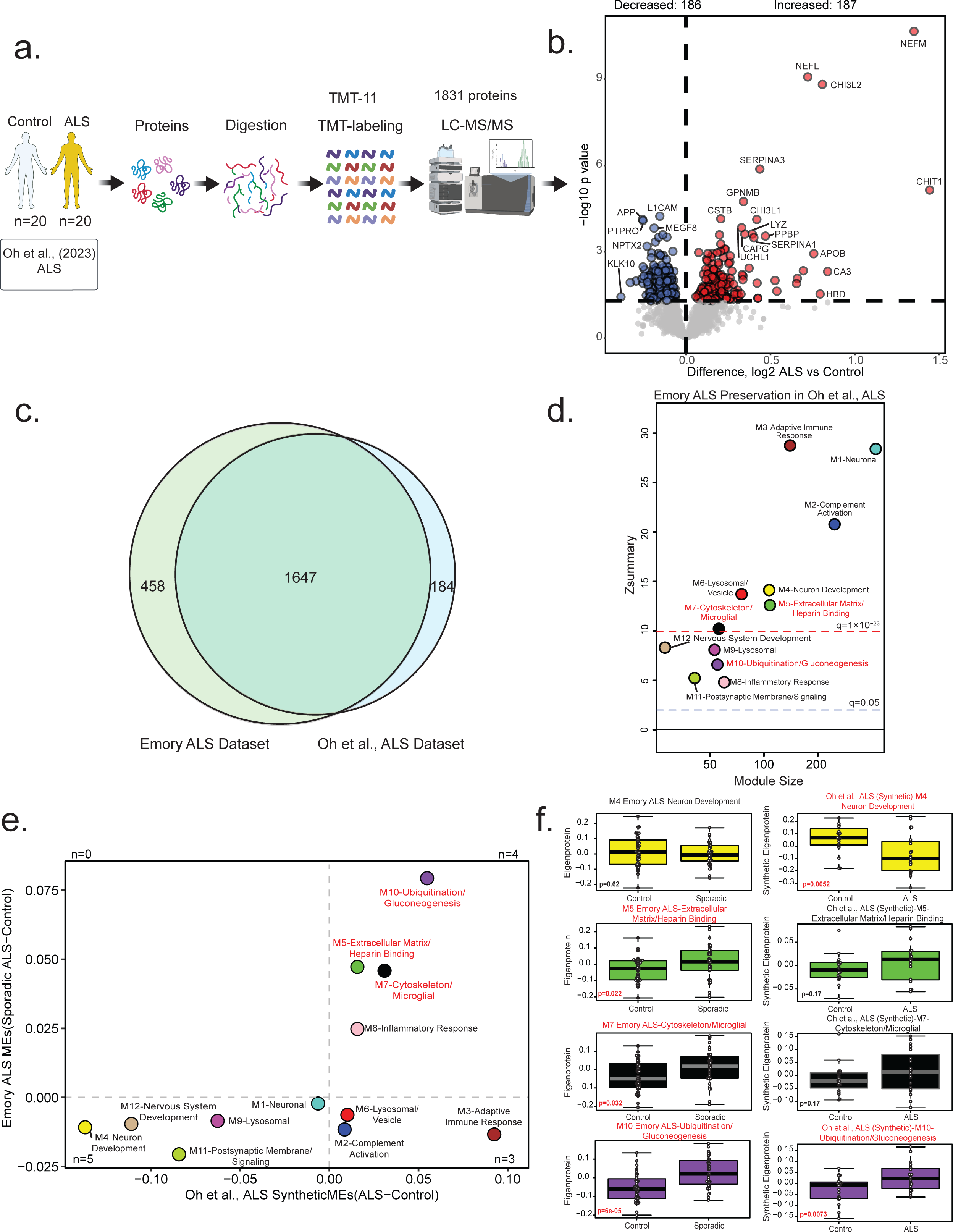
Emory ALS CSF protein network changes are preserved in an independent ALS CSF proteome. **a.** Schematic of experimental workflow to quantitatively evaluate similarities and differences across CSF proteomes generated from two different data sources (Emory ALS Set1 and Oh et al. ALS Set 2), indicating sample size, number of proteins quantified, and number of modules calculated. **b.** Volcano plot of Set 2 proteome indicating differential expression of ALS (n=20), versus control (n=20). **c.** Venn diagram representing the number of proteins that were quantified in the Emory Set1 and Set 2 ALS datasets. **d.** Module preservation of Emory Set 1 and Set 2 dataset. Number of proteins in each module (x-axis) is compared across Zsummary, and overall measurement of preservation, (y-axis). The red line at Zsummary=10 (q=1x10^-23^) indicates Emory ALS modules are highly preserved in the replication proteome. The blue line at Zsummary=2 (q=0.05) indicates Emory ALS modules are preserved in the independent replication dataset. **e.** Scatter plot representing differential abundance of Emory ALS module eigenprotein sALS – Control (y-axis) versus abundance of Oh et al. ALS synthetic eigenprotein ALS - Control (x-axis). **f.** Several paired Emory ALS module eigenprotein and Set 2 ALS synthetic module eigenprotein comparisons are presented. Synthetic eigenproteins were constructed for Emory ALS dataset and measured by disease type in Set 1 and Set 2 ALS datasets. A minimum of four proteins from the top 20% of module membership by kME (correlation to module eigenprotein) were used to assess synthetic eigenprotein value (y-axis) and compared across disease type (y-axis). Those that are significant (p<0.05) by one way ANOVA p-values are represented in red. Modules are identified by module number and top GO terms in Figure 3.

Our re-analysis of the independent Set 2 dataset revealed that the DAPs between ALS and control subjects closely resembled those presented in the publication (12) (**Figure 5b**). Of the 2,105 and 1,831 proteins identified from Set 1 and Set 2 proteomes, respectively, 78.2 and 90.0% of those proteins were shared (**Figure 5c**). The Emory ALS proteome (Set 1) included 458 proteins not found in the published Set 2 whereas Set 2 included 184 proteins not found in Set 1. All modules from the Emory Set 1 ALS dataset were found to be preserved in the Set 2 ALS proteome (**Figure 5d**). Modules associated with M1-Neuron, M2-Complement activation, M3- Adaptive immune response, M4-Neuron development, M5-Extracellular matrix/Heparin binding, M6-Lysosomal/Vesicle, and M7-Cytoskeleton/Microglia had Zsummary values over 10 (i.e. q=1x10^-23^) corresponding to being very well preserved. Modules associated with M9-Lysosome, M10-Ubiquitination/Gluconeogenesis, M11-Postsynaptic membrane/signaling, and M12- Nervous system development had Zsummary values over 2 (q=0.05) but less than 10, indicating preservation. Module eigenproteins and synthetic module eigenproteins (Emory ALS module first principal component in Set 2) were concordant in 9 modules, with three modules showing significant increase in Set 2 but a decrease in Set 1 (**Figure 5e**). Several module eigenprotein comparisons to synthetic eigenproteins were of particular interest, including the modules that were significantly different across disease and control in Set 1, as well as an additional module with M4-Neuron development ontology that was not significantly different in Set 1, but that was significantly different in Set 2 (**Figure 5f**).

Synthetic eigenproteins were compared across Set 1 and Set 2 CSF proteomes (**Supplemental Figure 1; Supplemental Table 8**). Synthetic eigenproteins associated with M1-Neuron, M2- Complement activation, M4-Neuron development, M11-Postsynaptic membrane/signaling, and M12-Nervous system development were found to be significantly different across disease (ALS versus Control) for the Set 2 proteomics. Thus, the two-dataset comparison recapitulates results, using case samples from different centers, with different TMT processing and analysis modalities. However, the lack of genomic traits in Set 2 prevents further validation of changes specific to C9orf72 expansion or SOD1 mutation.

### ALS-specific proteomic changes compared to Alzheimer’s Disease

To assess the specificity of the changes for ALS versus other neurodegenerative diseases, we performed analysis including TMT-MS proteomic data from 149 controls and 149 Alzheimer’s Disease (AD) previously reported (13). DAPs were compared across the ALS Set 1, ALS Set 2, and the Emory AD CSF datasets to determine which DAPs are associated with neurodegenerative disease in general and which proteins were associated with ALS or AD (**Figure 6**). Three different datasets from varying TMT types and different centers (Emory and Johns Hopkins (12)) (**Figure 6a**) were composed of a total of 220 disease cases plus 213 control individuals.

**Figure 6.**
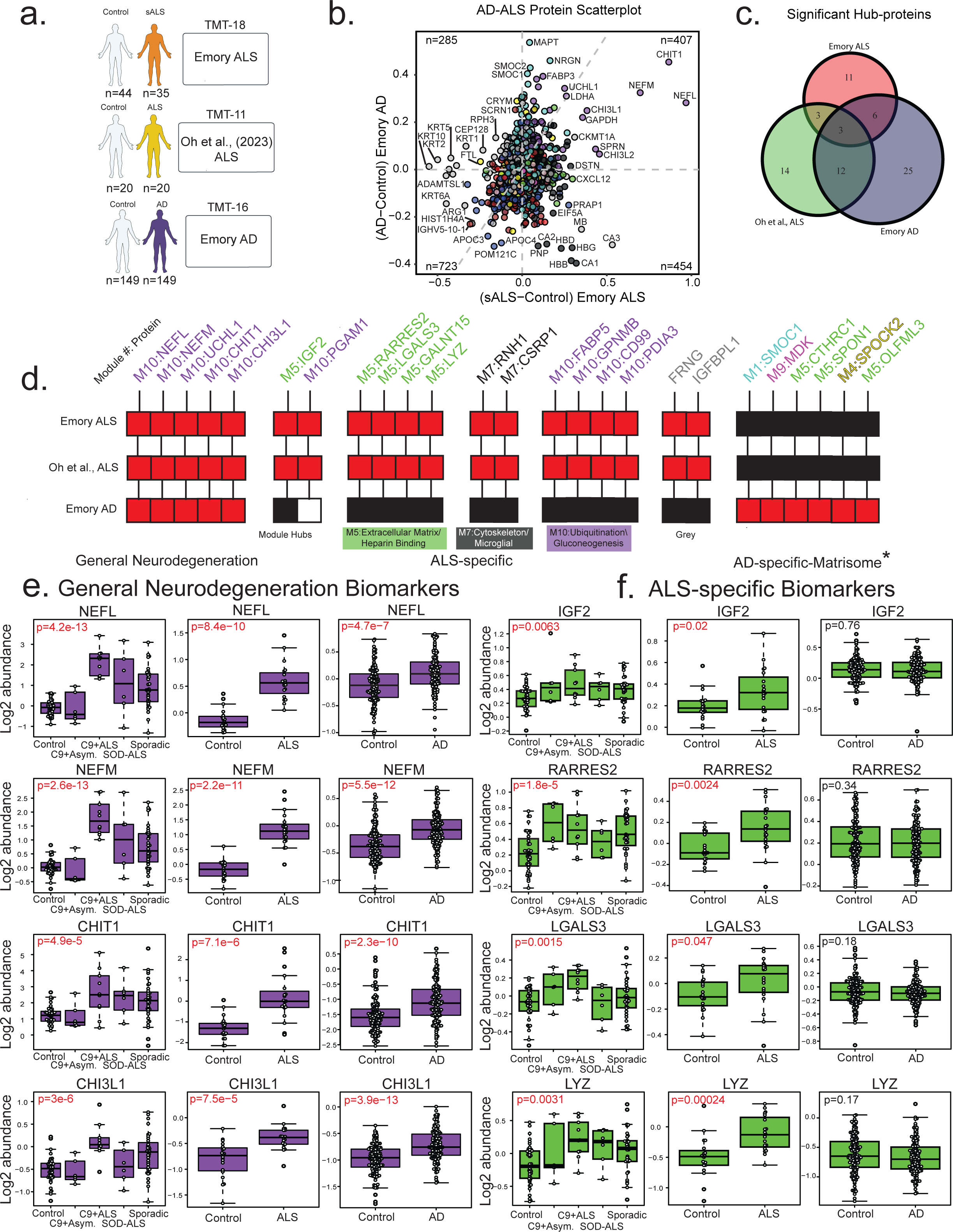
Assessing disease specificity of ALS CSF network and protein changes. **a.** Schematic of experimental workflow processing quantitative proteomics datasets with tandem-mass-tag (TMT) chemistry. **b.** Scatter plot indicating differential expression of sporadic ALS (n=35) versus the AD CSF dataset (AD; n=149). Log_2_ fold change of sporadic ALS (x-axis) and Log_2_ fold change of AD (y-axis) were compared for each protein (n=1,871). Proteins are colored corresponding to module. **c.** Venn diagram shows overlap of top ten module hub proteins, the proteins that most highly correlated with each module, (n=120) that significantly varied across disease type and control by one-way ANOVA. **d.** Heatmap indicating which datasets had proteins associated with disease both present and significantly different from controls (red), present and not significantly different from controls (black), or not present in the dataset (white). **e.** A selection of well characterized markers shared across neurodegenerative disease. For each marker the normalized, log_2_-transformed abundance (y-axis) and disease type (x-axis) are shown for each of the datasets in A, alongside p-values from one way ANOVA. **f.** ALS-specific changes, shared across ALS disease type and control but not AD.

Across Emory sALS-Control and Emory AD-Control 1,130 proteins were concordant, and 739 proteins were not changed within confidence limits (**Figure 6b**). The top 10 hub proteins were compared for significance across all three data sets with three found in common for all datasets and two specific to ALS (**Figure 6c**). Several proteins were found to be significantly different from controls in the same direction in all three proteomes including NEFL, NEFM, UCHL1, CHIT1, and CHI3L1, which can be linked to general neuroinflammatory or neurodegenerative processes occurring across diseases (**Figure 6d**). In total, two proteins from module hubs were significantly different from controls in the same direction in the ALS proteomes and were either not significantly different from controls or not detected in the disease specificity proteome (**Supplemental Table 9**). In addition, 14 proteins from modules representing M5-Extracellular matrix/Heparin binding, M7-Cytoskeleton/Microglia, and M10-Ubiquitination/Gluconeogenesis varied significantly, in the same direction, in ALS but did not vary significantly or varied significantly in the opposite direction in disease specificity. Of these proteins, 10 representative proteins are visualized (**Figure 6d**). Similarly, several proteins associated with AD were assessed in all three datasets including SMOC1, MDK, CTHRC1, SPON1, SPOCK2, and OLFML3. Four common neurodegenerative proteins NEFL, NEFM, CHIT1, and CHI3L1 were visualized to convey their significance across both ALS datasets and AD samples respectively (**Figure 6e**).

Four ALS-specific proteins IGF2, RARRES2, LGALS3, and LYZ were also visualized to demonstrate significance in ALS CSF proteomes but not AD (**Figure 6f**). These comparative analyses indicate that, in addition to protein markers of neurodegeneration consistent across ALS and AD, there are also differences associated with each disease phenotypes, which may be linked to underlying pathology. These differences may aid in elucidating disease mechanisms and identifying therapeutic targets for intervention.

## Discussion

In this study, we characterized the CSF proteome of several groups of ALS patients defined by genetic status and clinical diagnosis in order to identify shared and unique biomarkers across each ALS subgroup. By examining coordinated protein co-expression across genetic and sporadic forms of ALS, we identified shared and group-specific changes linked to gene ontology and cell-type function in the proteome. We compared sALS, C9orf72 ALS, C9orf72 asymptomatic carriers, SOD1 ALS, and healthy controls to identify biomarkers associated with each ALS type. Our analysis validated previously identified ALS associated CSF biomarkers (12, 38, 40, 41) but importantly, identified novel ALS biomarkers specific to genetic status as well as the onset of clinical symptoms.

Similar to previous studies, we found a significant increase in the abundance of neurofilament proteins (e.g., NEFL, NEFM) and chitinase associated proteins (e.g., CHIT1, CHI3L1, CHI3L2) across all clinically diagnosed ALS cohorts (12, 38, 40, 41). Increased abundance of neurofilament proteins in the CSF likely reflects the axonal degeneration of neuronal populations in the central nervous system (CNS) as these proteins are upregulated in the CSF of other neurodegenerative diseases including AD and frontotemporal lobar dementia (FTLD) (42, 43).

The functional consequence of increased abundance of CHIT1 and chitosan-like proteins (e.g., CHI3L1, CHI3L2) are unknown; these proteins have been shown to co-localize with markers of microglia and macrophages, thus they are likely associated with activation of microglia and/or macrophages (37). We further observed an increased abundance of other proteins associated with microglial activation, including GPNMB and LYZ. While increased GPNMB abundance has been previously reported in ALS CSF proteomes, the presence of LYZ has not been documented. However, both *GPNMB* and *LYZ* have been demonstrated to be upregulated at the transcript level in the spinal cord of ALS patients (9).

Unique to our study is the assessment of asymptomatic C9orf72 repeat expansion carriers. There is a strong age-related penetrance associated with C9orf72 ALS/FTD; thus, inclusion of asymptomatic C9orf72 carriers provides an early view into pre-symptomatic stages of disease (44, 45). Notably, established biomarkers discussed above that are commonly associated with neurodegeneration (NEFL and NEFM) and microglial activation (CHIT1, GPNMB, LYZ) were unchanged in asymptomatic C9orf72 carriers compared to controls. These data suggest that the observation of proteins associated with axonal degeneration and microglia activation in ALS cases are reflective of the presence of clinical symptoms. The increased abundance of protein markers thought to reflect neurodegeneration (NEFL and NEFM), only in the symptomatic C9orf72 carriers, provides internal validation of specificity of the proteomic findings. The identification of differences in protein abundance between C9orf72 carriers with and without ALS is particularly important, as it may identify prognostic markers of disease (such as AKR1B1, TKT, CFL1, CFL2, PPIA and FABP3). A shift to a pattern seen in symptomatic C9orf72 ALS could represent a precursor of imminent transition to symptomatic disease. It will be important to determine the relative order of changes prior to and during symptom onset in this genetically defined subset of individuals longitudinally, similar to recent analyses performed in autosomal dominant AD carriers, modeling protein trajectories relative to estimated year of onset (46).

Interestingly, Lactate dehydrogenase A chain (LDHA), which we observed to be upregulated in the CSF of symptomatic ALS cases and has previously been shown to be increased in the CSF of other neurodegenerative diseases, including AD (47), is also significantly increased in the CSF of asymptomatic C9orf72 carriers. LDHA is a glycolytic protein involved in the concomitant interconversion of pyruvate with lactate and NADH with NAD^+^(48). The robust increase of LDHA abundance in the CSF of both asymptomatic C9orf72 carriers and symptomatic ALS patients suggest that shifts in energy metabolism from oxidative phosphorylation to glycolysis (i.e. The Warburg effect (49, 50)) is an early cellular change associated with ALS prior to clinical onset. Alternately, lactate buffering between neurons and astrocytes is a contentious but well-described process potentially mitigating excitotoxicity (51, 52), and such excitotoxicity has been postulated to occur in ALS (53, 54).

Importantly, we identified several proteins that were robustly increased in the CSF of asymptomatic C9orf72 carriers but not C9orf72 ALS patients, including EMILIN1 and DNAJC3. Knockout experiments in mice and culture have shown that EMILIN1 plays a role in elastogenesis and maintenance of vascular cells (55). These findings were confirmed in bi-allelic, loss of function EMILIN1 families, demonstrating that not only does EMILIN1 assist EFEMP2 (fibulin-4, responsible for fibrillar collagen localization (56)) in elastogenesis, but also affects lysyl oxidase by EFEMP2 to establish collagen crosslinks (57). Similarly, an investigation into siblings that demonstrated loss of DNAJC3 has been shown to result in diabetes mellitus as well as “multisystemic neurodegeneration” (58). DNAJ/HSP40 proteins are also known to regulate HSP70 co-chaperones through stimulation of ATP hydrolysis (reviewed in (59)).

We also observed differential changes of neurodegenerative proteins in the CSF proteomes stratified by disease subgroup. For example, we found decreased levels of some keratins, similar to observations in CSF of asymptomatic AD versus AD (14). Intermediate filaments (like keratins and neurofilaments) have previously been implicated in toxic dipeptide repeats associated with C9orf72-associated ALS, with repeats promoting a high-density network of intermediate filaments in the cytoplasm (60, 61). This would suggest that dense networks of irregular intermediate filaments in the cytoplasm may have implications on phase separation of mutant TDP-43. Recently, our work has shown that keratins are significantly enriched in the cerebrovascular fraction of human brain and may be derived from smooth muscle cells (62), suggesting a role of cerebrovascular dysfunction in C9orf72-associated ALS potentially linked to TDP-43 pathology. In support of this hypothesis, keratins are not significantly decreased in abundance in SOD1 ALS, which does not have TDP-43 inclusions as a pathological hallmark.

Histone proteins (H1.5, H1e, H2A, H2A.Z, H2B type 2-F, and H4) were decreased in abundance relative to controls in genetic forms of ALS but not sporadic ALS, which may provide insight to differences in disease mechanisms. It may be that decreased abundance of histone proteins in genetic forms of ALS may be due to decreased histone deacetylases, which were not detected in CSF in our study. There are 18 histone deacetylases (HDACs) in humans that deacetylate histone and non-histone proteins (63). Decreased histone deacetylation has been linked to neurodegeneration (64–66) and HDAC abundance has downstream effects on HSP70 (67) as well as TDP-43 deacetylation and aggregation (68). Mutations in FUS (including the 521C mutation) cause a preferential interaction between FUS and RBM45, rather than HDAC1 (69). Histone acetyltransferases, deacetylases, and bromodomain (acetyl-lysine binding/reading domain- containing proteins) may present several druggable targets (70).

Though TDP-43 pathology is a neuropathological hallmark of sporadic and C9orf72-associated ALS, we did not identify TDP-43 in any CSF samples. Similarly, we did not identify cryptic peptides that are associated with TDP-43 dysfunction or the inclusion of cryptic exons (71, 72) despite including them in our proteomic databases for identification and quantification. Markers of TDP-43 pathology are not expected in SOD1-related ALS, however our systems biology approach identified a module (i.e., M8-Inflammatory response) as associated with SOD1, as well as other protein markers and modules. Total SOD1 protein was decreased in SOD1 ALS relative to controls and to the other patient groups. A recent study (73) found that mutant SOD1 was 16- fold lower in concentration in CSF than wildtype SOD1 and that the turnover of mutant SOD1 was two times faster (73). In conjunction with the finding that Tofersen lowered SOD1 by 30% (74), these results demonstrate the need to better understand how these concentrations are allocated amongst the myriad known SOD1 mutations and peptides. Our results therefore indicate that the underlying pathogenic differences between different genetic forms of ALS is reflected in the CSF proteome and is consistent with differences observed in the neuropathology and clinical course of these subgroups.

There are several shortcomings to our study that should be addressed in future analyses. Additional representation in ALS cohorts is necessary to confirm that biomarkers (such as NEFL, CHIT1, and recently elucidated novel biomarkers) are useful across diverse populations. Similarly, diverse cohorts may provide additional insight into ALS biology through a more complete view of how ALS affects patients. A larger sample size of genetic forms of ALS, the addition of asymptomatic cases, and the inclusion of other genetic forms of ALS (in particular cases associated with the RNA binding protein fused in sarcoma, FUS) would address further unmet needs. In addition, our dataset represents a snapshot of the biological processes at work in ALS. To this end, longitudinal clinical measures (10), combined with longitudinal proteomics may better help to better reveal these processes. The identification of additional ALS biomarkers will inform future efforts to identify disease onset and track ALS progression.

## Conclusion

Our investigation identified protein biomarkers that may distinguish the transition from asymptomatic to symptomatic phase and differences between genetic and sporadic forms that indicate differences in mechanisms of disease. We incorporated different external and disease-specific datasets to validate our proteome and distinguish ALS-specific pathways. Future analyses should utilize these differences to determine their applicability to diverse ALS cohorts and identify how these biomarkers develop longitudinally.

## Availability of Data and Materials

Raw mass spectrometry data and pre- and post-processed CSF protein abundance data and case metadata related to this manuscript are available at https://www.synapse.org/EmoryALS.

---

**Supplemental Figure 1.** Synthetic eigenproteins were constructed for Emory ALS dataset and measured by disease type in Emory ALS dataset (Set 1) and the replications ALS dataset (Set 2). A minimum of four proteins from the top 20% of module membership by kME (correlation to module eigenprotein) were used to assess synthetic eigenprotein value (y-axis) and compared across disease type (y-axis). Where significant (p<0.05) by one way ANOVA p-values are represented in red. Modules are identified by module number and top GO terms.

